# The role of dopaminergic, cholinergic and noradrenergic networks in hyposmia in Parkinson’s disease

**DOI:** 10.1101/2024.10.23.619791

**Authors:** Jean-Baptiste Pérot, Esther Kozlowski, Audrey Fraysse, Salim Ouarab, Romain Valabregue, Sana Rebbah, François-Xavier Lejeune, Emma Massy, Rahul Gaurav, Isabelle Arnulf, Jean-Christophe Corvol, Marie Vidailhet, Nadya Pyatigorskaya, Cécile Galléa, Stéphane Lehéricy

## Abstract

**Background:** Olfactory impairment is frequently observed in Parkinson’s disease (PD) before motor symptom onset. Patients with isolated rapid eye movement sleep behaviour disorder (iRBD) are at risk to develop PD and present olfactory dysfunction. Subcortical neuromodulatory nuclei including the substantia nigra, the nucleus basalis of Meynert, and the locus coeruleus, may all contribute to olfactory dysfunction.

**Objective:** The objective of this study was to understand the network alterations underlying olfactory dysfunction in PD and in the early prodromal iRBD patients using multimodal MRI.

**Methods:** PD (n = 107), iRBD patients (n = 35) and healthy controls (HC, n = 39) were recruited from the ICEBERG cohort for a cross-sectional study. We separated subjects in Healthy Controls, iRBD patients, and PD patients with or without RBD and with or without anosmia. Olfactory, motor, and cognitive scores were assessed and combined with multimodal imaging.

**Results:** We found that olfaction positively correlated with (i) striatal DaT signal in PD patients; (ii) neuromelanin contrast in the locus coeruleus and (iii) Nucleus Basalis of Meynert grey matter (GM) volume in all patients. These signals were uncorrelated with motor and cognitive scores. Functional connectivity was reduced in regions of the cholinergic olfactory network in anosmic patients. Functional connectivity was also reduced in the noradrenergic network of patients with RBD.

**Discussion:** Our results indicate the implication of the cholinergic network in PD patients with anosmia and a contribution of the noradrenergic network to olfactory dysfunction, only in patients with RBD.

## Introduction

Although motor manifestations are the central clinical feature of Parkinson’s disease (PD), non-motor features are observed in most patients with PD. Olfactory dysfunction is one of the most frequent non-motor manifestations and is considered a supportive criterion for PD^1^. Hyposmia occurs early and often predates the onset of motor symptoms. Olfactory deficit is thus frequently observed in prodromal PD conditions such as isolated rapid eye movement sleep behavior disorders (iRBD)^2^.

Processing of olfactory information engages a large network, with an organized system of connections. The olfactory bulb, receiving projections from the olfactory epithelium, is connected to cortical and subcortical structures^3^. Structures of the olfactory system include the anterior olfactory nucleus, the entorhinal cortex, the piriform cortex, the olfactory tubercle, the amygdala and peri-amygdaloid cortex, the hippocampus and the orbitofrontal and insular cortices. Subcortical neuromodulatory nuclei are also connected with the olfactory structures and the olfactory bulb including the noradrenergic locus coeruleus/subcoeruleus (LC), the cholinergic basal forebrain nuclei, such as the Nucleus Basalis of Meynert (NBM) and the dopaminergic substantia nigra (SN)^4^. Numerous imaging studies have indeed shown that olfactory impairment in PD was associated with reduced olfactory bulb volume and increased depth of the olfactory sulcus^5–9^, cortical gray matter abnormalities in olfactory regions^10–14^ as well as impairment of subcortical cholinergic^15^ and dopaminergic circuits^15–18^. Few studies have investigated prodromal PD with variable results^12,19,20^. However, no study has investigated the relationship between hyposmia and the impairment of these different structures together and their link with cognition in a large number of subjects with PD and iRBD.

Olfactory damage is also important to study because the olfactory bulb could be a predominant entry route for PD pathology^21^. PD is a synucleinopathy characterized by the presence of abnormal intraneuronal misfolded α-synuclein deposition. Studies have suggested that these aggregates may propagate from cell-to-cell in a prion-like manner^22^ with two possible entry routes from the enteric peripheral nervous system to the brainstem and from the olfactory bulb ^21^. Imaging studies have recently provided arguments in favor of this hypothesis suggesting that the propagation of PD-related pathology could be divided in two main profiles^23,24^ called body-first and brain-first profiles^23^. The body-first (bottom-up) profile corresponds to a propagation route starting in the enteric or peripheral autonomic system and spreading to the brain through brainstem nuclei such as the dorsal motor nucleus of the vagus nerve. The brain first (top-down) profile is associated with initial pathology in the brain, probably originating from the olfactory system, progressively spreading to the lower brainstem nuclei. The presence or absence of RBD appeared to be a determining factor of one or the other model. The model that best explained the imaging data observed in PD patients with RBD and iRBD was the “body first” model while the one that best explained the data observed in PD patients without RBD was the “brain first” model^23–25^. However, these two models do not fully account for the heterogeneity of the disease. For instance, the high prevalence and severity of olfactory dysfunction in iRBD patients and the fact that cognitive and olfactory disorders are frequent and early symptoms in PD patients with RBD is not explained by the body-first model. Other routes and greater individual variability than these two trajectories may thus be observed in PD that need to be explored.

To address these questions, we studied a cohort of 107 PD patients with and without RBD, 35 patients with iRBD and 39 healthy controls from the ICEBERG study evaluated using extensive clinical, polysomnography, multimodal imaging examinations including multimodal MRI with neuromelanin-sensitive, structural and diffusion imaging and dopamine transporter SPECT. The objectives of this study were to identify the cortical and subcortical networks underlying olfactory dysfunction in PD and to evaluate the involvement of dopaminergic, cholinergic and noradrenergic networks in patients with RBD (PD and iRBD). We tested the hypotheses that the olfactory disorder would be associated not only with damage to the cortical olfactory regions but also to the subcortical neuromodulatory circuits (SN, LC, NBM) and that this damage would also be important in patients with RBD.

## Materials and Methods

### Participants

Participants were prospectively recruited from November 2014 to February 2020 as part of the ICEBERG longitudinal cohort (ClinicalTrials.gov: NCT02305147). This cohort included healthy control subjects, patients with iRBD, and patients with PD. The inclusion criteria for PD comprised an age between 18 and 75 years, a diagnosis of PD meeting the Movement Disorder Society PD criteria^1^, no or minimal cognitive disturbances (defined as a Mini-Mental State Examination [MMSE] score greater or equal to 26/30), and a disease course lower than 4 years, defined as the period from first manifestation of a cardinal motor parkinsonism (as reported by the participant), to inclusion time. The usual treatment of participants was not stopped during evaluations. Patients with iRBD had a history of dream-enacting behaviours with (potentially) injurious movements and enhanced tonic chin muscle tone or complex behaviours during REM sleep but did not meet the criteria for Parkinson’s disease or dementia^26^. The inclusion criteria for healthy participants comprised the absence of present or past neurological or psychiatric disorder (as assessed after interview and clinical examination by a neurologist; note that sleep disorders were not a reason for exclusion in PD and control groups) and being matched for age and sex with participants with PD. All participants signed a written informed consent to take part in the study, which had been approved by the local ethics committee (CPP-Ile de France-Paris 6, RCB 2014-A00725-42).

### Clinical evaluation

All participants underwent clinical examinations to assess motor, olfactory, and cognitive abilities at the time of MRI recordings. The Movement Disorders Society - Unified Parkinson’s Disease Rating Scale part III with no dopaminergic treatment in the last 12h (MDS-UPDRS-III) assessed the severity of motor symptoms^27^. The Pennsylvania Odour Identification Score (UPSIT) to assessed olfactory function based on the recognition of common odours. Forty odours were consecutively presented with a “a scratch and smell” panel. Participants identified the smell from four options, without time constrain. The Mattis Dementia Rating Scale^28^ assessed cognitive symptoms.

Groups of participants were defined based on three criteria: the UPSIT score, the presence of RBD, and the presence of PD. We considered patients with anosmia when they had an UPSIT score strictly inferior to 25^29^. The presence of PD was assessed during the clinical examination at inclusion. The presence of RBD was confirmed by polysomnography in all participants. Five groups were defined including healthy controls (*HC)*, patients with iRBD (*iRBD*), patients with PD, RBD and anosmia 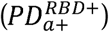 , patients with PD and anosmia without RBD 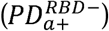 , and patients with PD but no RBD and no anosmia 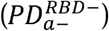 .

### Imaging protocol

MRI was performed for all participants using a Siemens Prisma 3T scanner (Siemens Healthineers) with a 64-channel head reception coil. The protocol included acquisition of a whole brain anatomic image, NM-sensitive image covering the SN and LC, and whole brain resting-state fMRI. DaT single-photon emission tomography was performed on a subset of patients and health controls using hybrid gamma camera Discovery 670 Pro system (GE Healthcare) with ^123^I-N-omega-fluoropropyl-2beta-carbomethoxy-3beta-(4-iodophenyl)nortropane (^123^I-FP-CIT) tracer (DaTScan^TM^). The detailed imaging protocols for MRI and DaTScan are presented in Supplementary Information.

### Image analysis

#### Image registration

Whole brain T1-weighted images were registered to Montreal Neurological Institute template for Voxel-based morphometry analysis with SPM12^30–32^.

NM-sensitive images were coregistered to the anatomical images using affine transformation.

DaTScan^TM^ images were coregistered to the MP2RAGE image using rigid transformation and partial volume correction^33^. For each subject, the DaT striatal binding ratio (DaT-SBR) was the signal intensity in the region of interest (ROI) normalized to a reference region in the occipital lobe. The reference region was segmented by co-registering the automated anatomical labelling template to the anatomical image^34^. The average DaT-SBR was then extracted in the putamen using the ROI of the YeB atlas^35,36^ that was individually segmented on the anatomical T1 image.

#### Segmentation of olfactory network and subcortical nuclei

The SN and LC were segmented on the NM-sensitive image as detailed in Supplementary Information.

ROIs belonging to the olfactory network were isolated from the FreeSurfer segmentation. ROIs were separated in two subtypes: cortical olfactory areas and subcortical olfactory areas which were used for VBM analysis and are detailed in Supplementary Information.

#### Functional image preprocessing

Functional EPI data were processed using standard tool proposed by SPM12 (REF). In short, we perform slice-timing correction, motion correction, and coregistration to the T1w volume. Finally the EPI volume were resliced at 1.5 mm isotropic in the Montreal Neurological Institute (MNI) space using non-linear deformation computed on the T1w volume.

#### React analysis

Receptor-enriched analysis of functional connectivity was performed following the approach described in ^37^. PET atlas of cholinergic and noradrenergic receptors was used as spatial regressors for functional connectivity computation of rs-fMRI data. Fsl tools were used to fit this spatial GLM in order to obtain representative time series for each receptor A mask of the NBM and LC were respectively used for cholinergic and noradrenergic receptors enriched analysis of functional connectivity. The resulting time series were used (with SPM) as regressors in a second multiple regression of BOLD data to estimate maps of functional connectivity corresponding to activation of cholinergic or noradrenergic receptors of the NBM or LC.

### Statistical analyses

All statistical analyses for clinical, NM-CNR, and DaT metrics were performed using JASP (JASP Team, 2023, Version 0.17.3). Groups were compared using Analysis of Covariance (ANCOVA) with age as a nuisance factor and Tukey’s post-hoc analysis. Correlations with UPSIT scores were tested in all patients using Pearson’s correlation with no hypothesis on normality.

VBM analysis was performed on individual VBM grey matter volume maps (obtained after segmentation of the T1w anatomical images) with SPM12 adjusted considering age, sex, and TIV. These adjusted values were entered in a one-way ANOVA model with five levels (*HC*, *iRBD*, 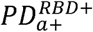 , 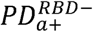 , 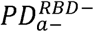). We evaluated the group differences and the direction of the group differences by defining the main effect and post-hoc t-tests. We tested the correlation between adjusted values and UPSIT considering all patients (*iRBD*, 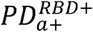 , 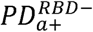 , 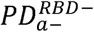). Family-wise error for multiple comparisons was applied with *p*<0.05. For both group comparisons and correlations, statistical analyses were performed separately on the cortical and subcortical subparts of the olfactory network.

### Data availability

Data is available to download in supporting data.

## Results

### Population

#### Demographics

Four groups were matched for age (*HC*, *iRBD*, 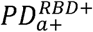 and 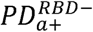) but 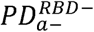 were significantly younger. PD patients were in the early phase of the disease with a mean disease duration of 1.6±1.1 years. Except for *iRBD* that was more frequent in men, groups were matched for sex. Age and sex were thus considered as covariables of non-interest in all statistical analyses. As expected, groups with anosmia had lower UPSIT scores than groups without anosmia. Clinical data of each group is presented on **Table 1**.

**Table 1:**
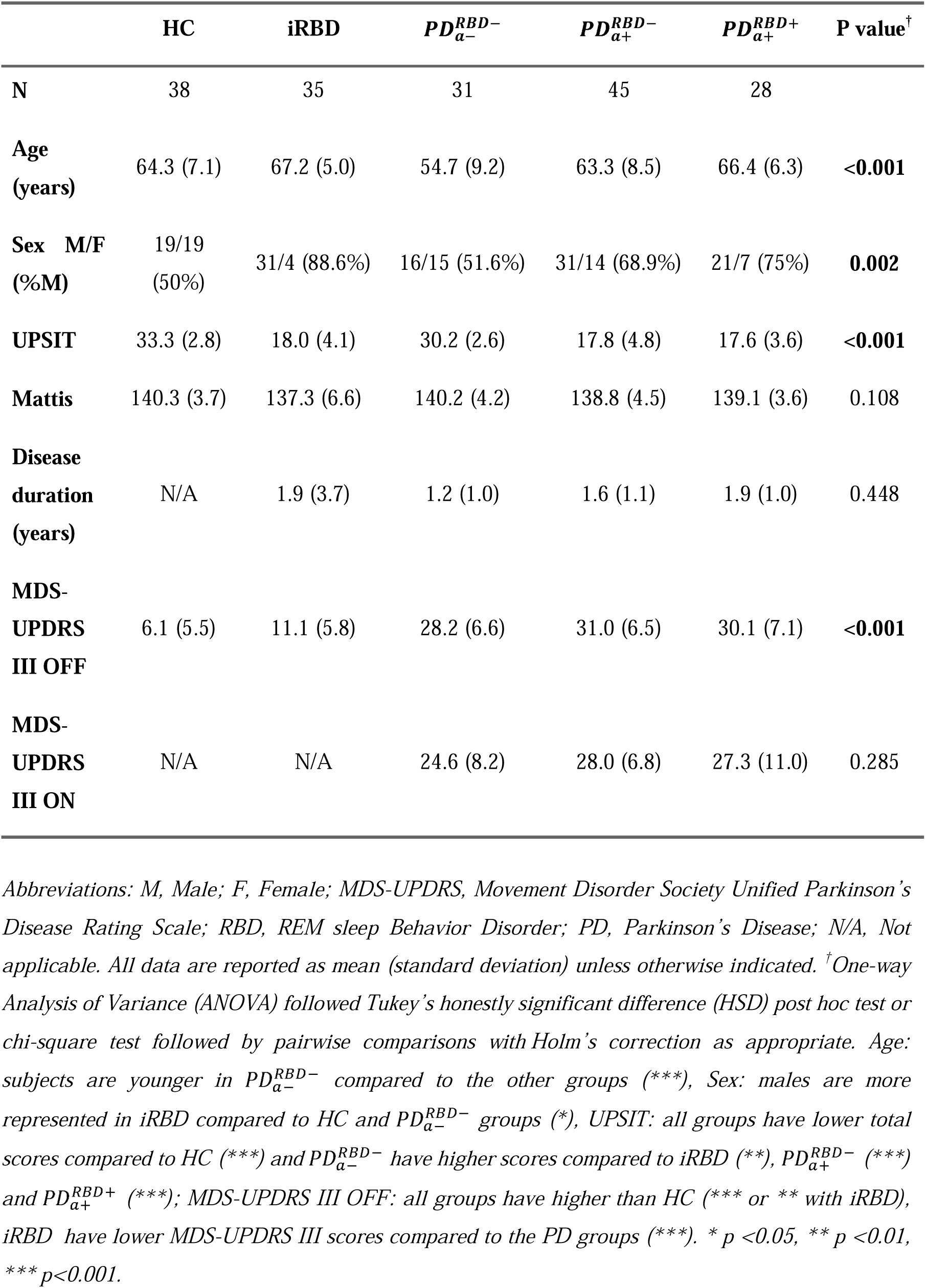
Summary of clinical data.

None of the groups showed significant cognitive change as measured with the Mattis score. MDS-UPDRS III scores were significantly different between groups (ANCOVA, F_4_=127.9), increased in *iRBD* compared with HC (Tukey, *p*=0.014), and in PD patients compared with *iRBD* (*p*<0.001) and *HC* (*p*<0.001). MDS-UPDRS III scores were similar between PD groups. UPSIT scores differed between groups (F_4_=136.6), were as expected reduced in all anosmic patients (*iRBD*, 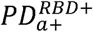 and 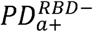 , all *p*<0.001) and did not differ between these groups. 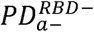 group showed reduced UPSIT compared with HC (*p*=0.003), but superior to anosmic groups (*p*<0.001). **Supplementary Figure 1** presents the distribution of UPSIT, Mattis, and tMDS-UPDRS III scores in all groups.

#### Correlations between UPSIT and clinical scores

UPSIT scores significantly correlated with age when considering all patients (Pearson’s correlation, *p*<0.001), while no correlation was found in *HC* (**Supplementary Figure 2**). In anosmic subjects only (*iRDB*, 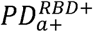 , 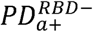), the correlation between UPSIT scores and age was no longer significant. UPSIT scores did not correlate with Mattis or MDS-UPDRS III scores, whether in all patients or in anosmic patients only.

### Alterations of noradrenergic and dopaminergic nuclei are observed using Neuromelanin-MRI and DaTscan

#### Group effect

There was a significant effect of the group on the NM-CNR in the LC (ANCOVA, F_4_=4.8) and SN (F_4_=12.9), as well as on the putaminal DaT-SBR (F_4_=120.2). In the LC, NM-CNR was significantly decreased in *iRBD* and 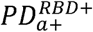 patients compared to HC (Tukey, *p*=0.004, **Supplementary figure 3a**) and 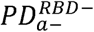 (*p*=0.003); but remained unchanged in the other groups without RBD. In the SN, NM-CNR was decreased in all anosmic patients compared with HC (*p*<0.001, **Supplementary figure 3b**), and in 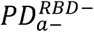 patients (*p*=0.044). Significantly greater severity was found in 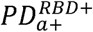 patients compared to *iRBD* (*p*=0.027) and 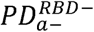 (*p*=0.006). In the putamen, DaT-SBR was reduced in all PD patients compared with HC and *iRBD* (*p*<0.001, **Supplementary figure 3c**), as well as in *iRBD* patients compared with HC (*p*=0.001). DaT-SBR did not differ between the PD groups.

#### Correlations with UPSIT scores

In all patients, UPSIT scores correlated with NM-CNR in the LC (**Figure 1**, *p*<0.01) but not with NM-CNR or volume in the SN or DaT-SBR. NM-CNR in the LC also correlated with NM volume in the SN (**Supplementary Figure 4**, *p*<0.05). MDS-UPDRS-III scores negatively correlated with DaT-SBR in all subjects including *HC* (**Supplementary Figure 4**, *p*<0.001) but not with NM markers in the SN. UPSIT scores did not correlate with cognitive (Mattis) or motor (MDS-UPDRS-III) scores.

**Figure 1:**
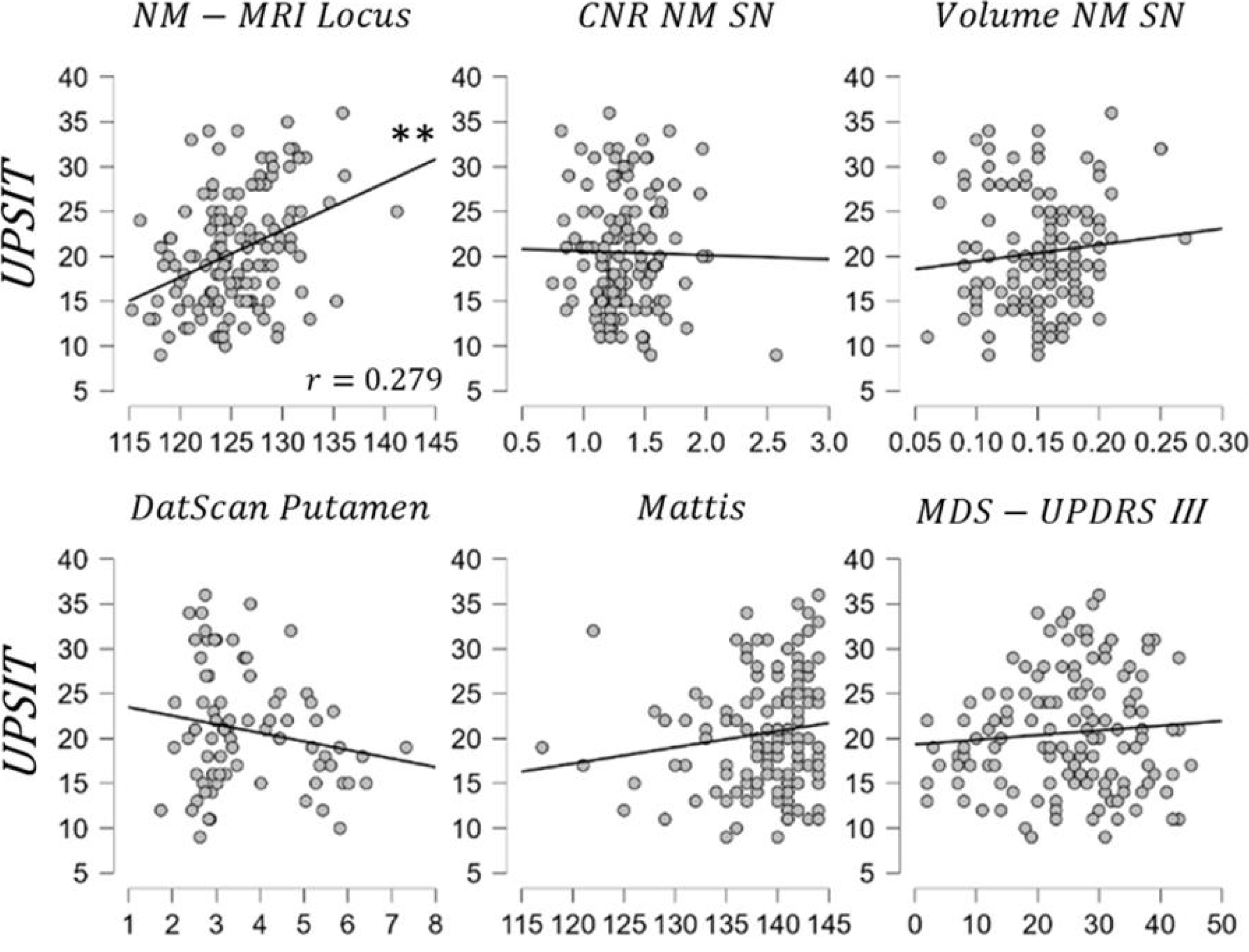
Scatter plots with linear regression lines of UPSIT scores versus other markers in all patients. Pearson’s R correlation coefficients. Significant correlation was only observed between UPSIT scores and NM-CNR in the LC and is indicated using stars (** indicate p<0.01).

### Atrophy of cholinergic nucleus is shown by Voxel-Based Morphometry

We evaluated the impact of anosmia associated with RBD and PD on grey matter volume in the brain using VBM. There was no significant difference in grey matter volume in any patient groups compared to *HC*.

We investigated the correlation between individual UPSIT scores and grey matter volume in all patients considering the cortical and subcortical olfactory masks separately.

In the cortical mask, we found significant correlations between UPSIT and grey matter volume in the insula, amygdala, anterior cingulate cortex, parahippocampal cortex, orbitofrontal cortex, and primary sensory cortex (**Figure 2)**. In the subcortical mask of the olfactory network, correlations between VBM and UPSIT were found in the NBM and in cerebellum (**Figure 2**, R=0.39, *p*<0.001).

**Figure 2:**
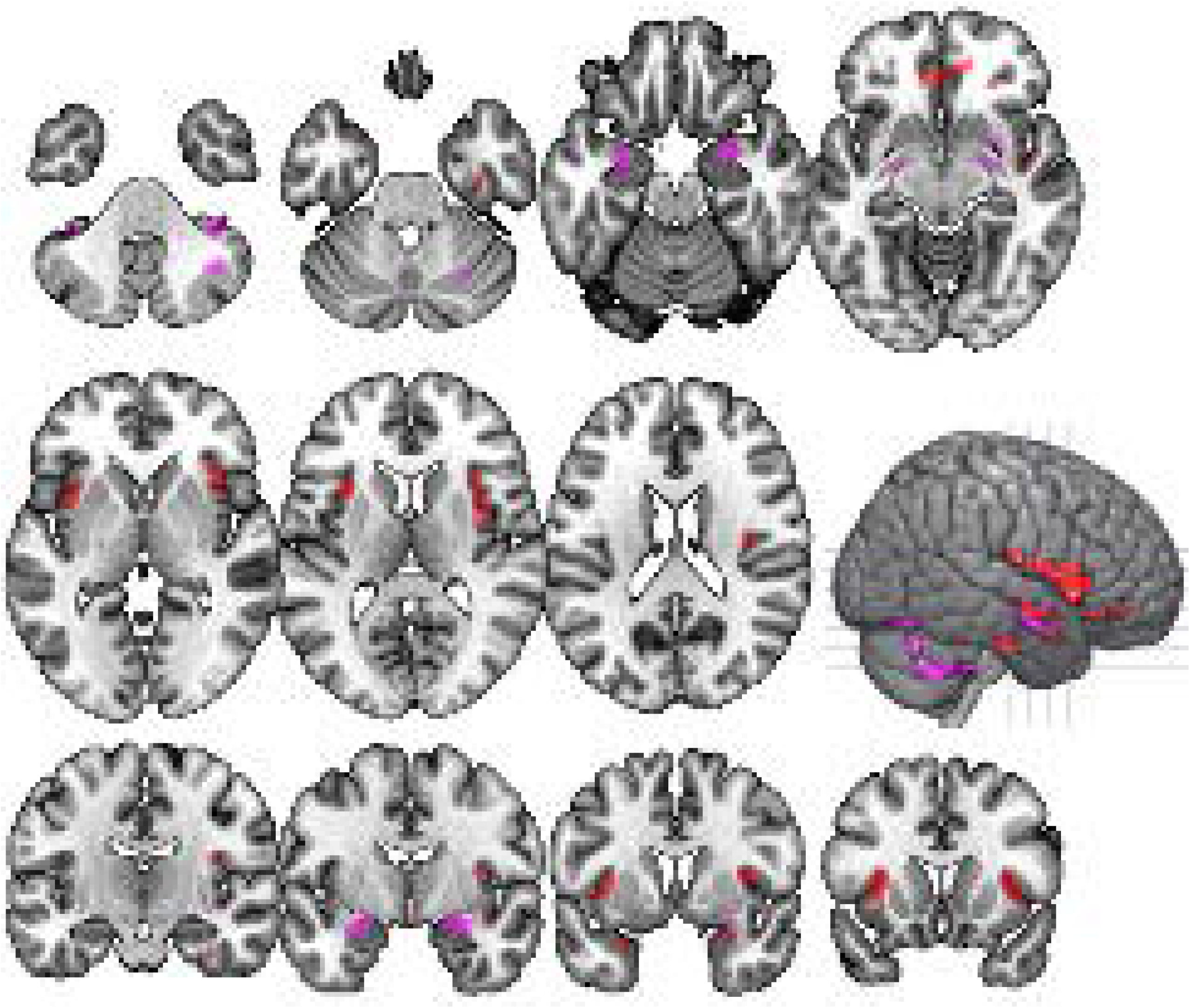
Correlations between UPSIT and VBM analysis in the cortical olfactory areas (red) and in subcortical olfactory areas (purple). Pearson’s correlation adjusted with age and sex. Right is right.

### Damage to dopaminergic, noradrenergic and cholinergic nuclei in relation to hyposmia and RBD

To study the contribution of subcortical noradrenergic, dopaminergic and cholinergic neuromodulatory nuclei in each disease group, we examined imaging markers in the three regions of interest of the LC (NM-CNR), putamen (Dat-SBR) and NBM (grey matter volume), respectively. **Figure 3** plots individual data in 3D along three axes, namely the putaminal SBR, the NM-CNR in the LC, and grey matter volume in the NBM. We represented the effect of PD on **figure 3A**. All PD patients were grouped and can be identified due to reduced DaT-SBR compared with *HC* and *iRBD* patients. **Figure 3B** presented the effect of RBD by grouping *iRBD* and 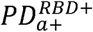 patients on the one hand, and 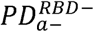 and 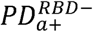 patients on the other hand. The two groups can be separated using NM-CNR in the LC. Finally, on **figure 3C** is shown the effect of anosmia by grouping *iRBD*, 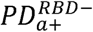 and 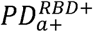 . Anosmic patients can be distinguished from *HC* and 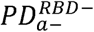 due to reduction of both NM-CNR in the LC and NBM grey matter volume.

**Figure 3:**
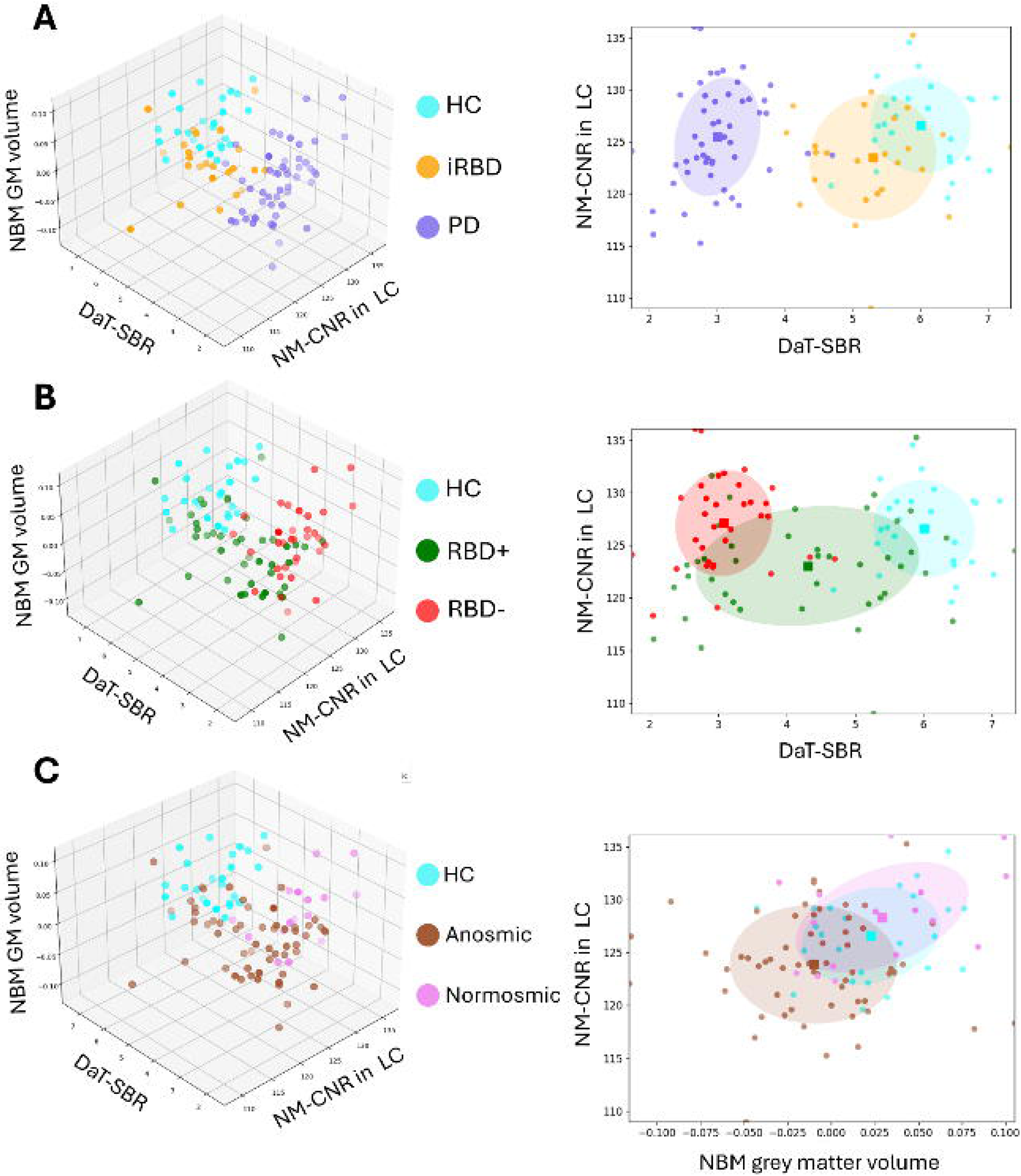
Differences between groups in subcortical neurotransmitter systems. Left: 3D scatter plot of all subjects as a function of Putaminal SBR, NM contrast in the LC, and NBM grey matter volume. Right: 2D projection with contour plot representing covariance. A) Effect of PD. Subjects were grouped by HC, iRBD and PD patients (including 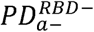 , 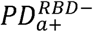, *and* 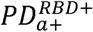 ). B) Effect of RBD. Subjects were grouped by HC, patients with RBD (including *iRBD* and 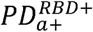) and patients without RBD (including 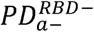 and 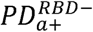). C) Effect of anosmia. Subjects were grouped by *HC,* anosmic (including *iRBD,* 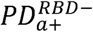 *and* 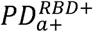) and normosmic 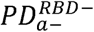 ).

### Cholinergic and noradrenergic functional connectivity is decreased in anosmic patients

To further investigate the role of neurotransmitters in olfactory dysfunction in PD patients with and without RBD, we used functional connectivity analysis enriched with neurotransmitter receptor maps (REACT analysis). We focused on cholinergic and noradrenergic receptors because the NBM and LC changes correlated with UPSIT scores, and did not include dopaminergic receptors that did not.

Significant differences in functional connectivity were found between HC and the patient groups with anosmia. However, no significant correlation was found between UPSIT scores and functional connectivity enriched with the cholinergic and noradrenergic neurotransmitters.

#### NBM cholinergic receptors-enriched functional analysis

Cholinergic receptors-enriched functional analysis showed decreased functional connectivity between the NBM and other brain areas in the anosmic groups only (**Figure 4A**). In patients with RBD compared with *HC*, decreased functional connectivity was found between the NBM and the frontal cortex including the left dorsolateral frontal cortex in *iRBD* (**Figure 4A**, green) and bilateral orbital and ventrolateral and dorsolateral frontal cortex in 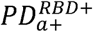 (**Figure 4A**, yellow). In 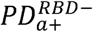 , decreased functional connectivity was found between the NBM and the left anterior putamen (**Figure 4A**, purple). No clusters were found in 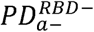.

**Figure 4:**
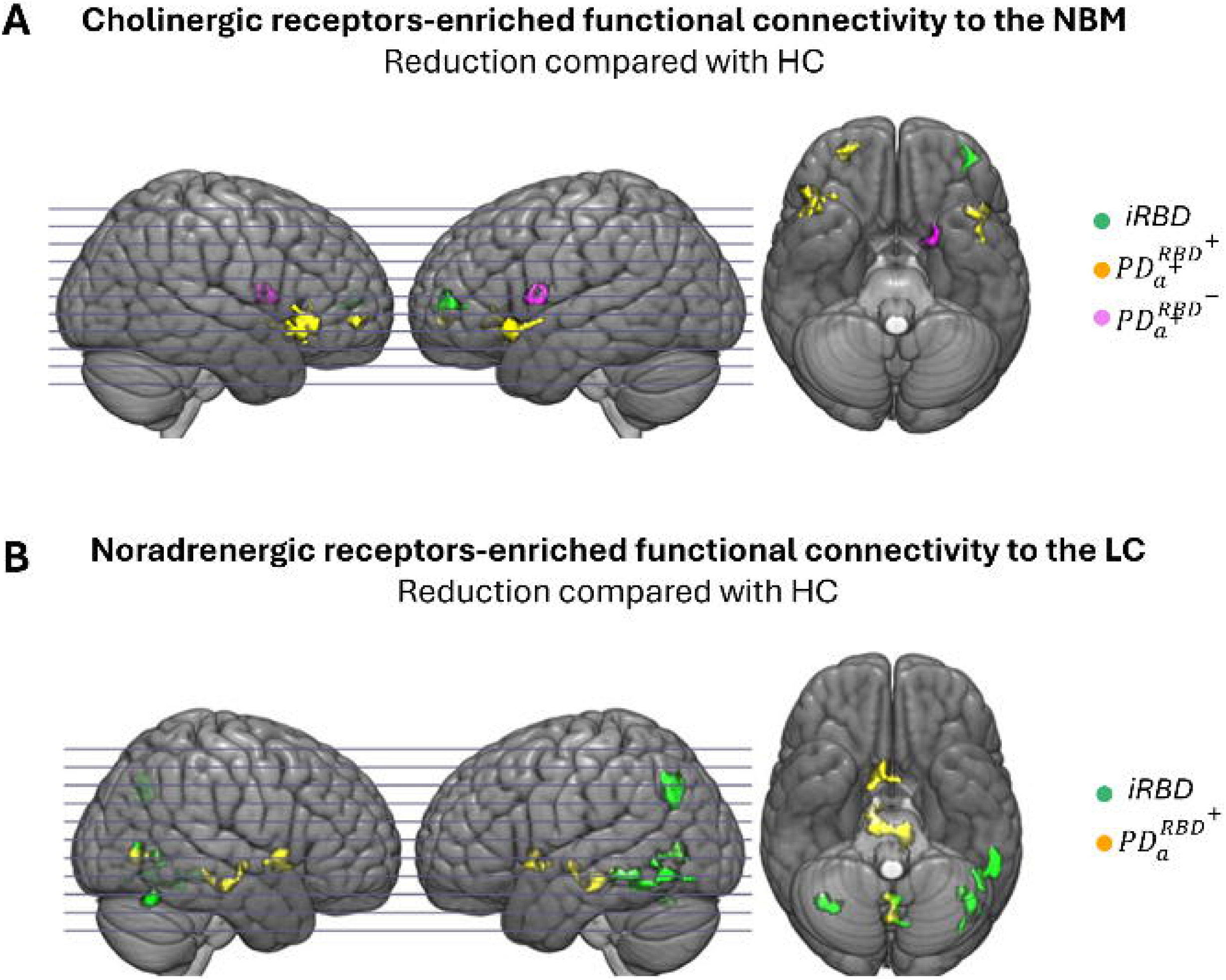
Receptors-enriched Analysis of Functional Connectivity (REACT). A) Regions with reduced cholinergic receptors-enriched functional connectivity to the NBM in iRBD (green), 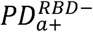 (violet), and 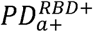 (yellow) compared to *HC.* B) Regions with reduced noradrenergic receptors-enriched functional connectivity to the LC in iRBD (green), 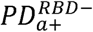 (purple), and 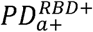 (yellow) compared to *HC.* Right is right on axial views.

#### LC noradrenergic receptors-enriched functional analysis

Noradrenergic receptors-enriched functional analysis showed decreased functional connectivity between the LC and other brain areas in patients with RBD (*iRBD* and 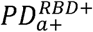) but not without RBD (**Figure 4B**). In *iRBD*, clusters were found the left parietotemporal areas and the right inferior occipitotemporal cortex. In 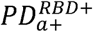 , clusters were found in the right SN and caudate nucleus and left tegmental areas. No changes were found in PD patients without RBD ( 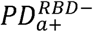 and 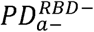).

## Discussion

We found that reduced performance in the olfactory test correlated not only with markers of atrophy in a cortical network involving the olfactory regions, in particular the amygdala and the orbitofrontal cortex, but also with damage to the subcortical cholinergic (grey matter in the nucleus basalis) and noradrenergic (neuromelanin locus signal) nuclei. In contrast, the olfactory deficit was not correlated with the impairment of striatal dopaminergic function (DAT) or imaging markers in the substantia nigra (neuromelanin signal). The olfactory deficit was independent of the motor and cognitive deficits. In addition, changes in grey matter in the NBM and reduced cholinergic receptors-enriched functional connectivity of the NBM with the cortex were observed in all anosmic patients whereas changes in neuromelanin signal in the LC and reduced noradrenergic receptors-enriched functional connectivity of the LC with the cortex appeared mostly associated with the RBD status.

### Cortical alterations related to olfactory dysfunction in PD

Olfactory dysfunction in PD and iRBD was associated with cortical atrophy of many regions of the olfactory network including the insula, amygdala, anterior cingulate cortex, parahippocampal cortex, and orbitofrontal cortex. These results highlight the vulnerability of this network in PD and its association with anosmia in line with previous studies that reported changes in grey matter^10,11,14,38^ or in diffusion measurements^39–41^ in olfactory regions in PD. Reduced olfactory bulb volume and changes in olfactory sulcus were also reported in these patients, structures that were not studied here^5–9^. In contrast, patients had no detectable change in overall grey matter volume compared to HC, which is explained by the early phenotype of patients with a mean disease duration of 1.6 years. Lastly, cortical atrophy did not correlate with cognitive scores in the same regions, suggesting that the observed correlation was specific to anosmia.

### Subcortical alterations related to olfactory dysfunction in PD

#### Contribution of the cholinergic pathway to olfactory dysfunction in PD

UPSIT scores also correlated with alterations in subcortical cholinergic and noradrenergic nuclei that were thus associated with olfaction deficits. Reduced grey matter in the cholinergic NBM correlated with the decrease in UPSIT scores. Reduced cholinergic receptors-enriched functional connectivity of the NBM with several frontal cortical regions including the orbitofrontal cortex was observed in all anosmic patients but not in normosmic patients. Histological studies have reported early degeneration of cholinergic neurons of the NBM in PD^42–44^. Previous PET study using acetylcholinesterase tracer showed that cholinergic denervation of the limbic cortex was a robust determinant of hyposmia in subjects with moderately severe PD^15^. Another study using MRI reported a correlation between reduced grey matter in the basal forebrain and UPSIT scores^13^. Damage to the NBM is frequently observed in neurodegenerative diseases with olfactory dysfunction such as Alzheimer’s disease^29^. Cholinergic neurons of the basal forebrain provide the innervation of the cortex including the olfactory regions, with cholinergic neurons of the horizontal limb of the diagonal band projecting to the olfactory bulb and of the NBM to the cortex and the amygdala^45^. The NBM has thus been associated with cognitive deficits in PD using structural MRI^46,47^, functional MRI^48^ and PET with cholinergic tracers^15^. In the present study, patients were in the early phase of the disease and did not present cognitive deficits. Moreover, grey matter changes in the NBM correlated specifically with olfactory and not with cognitive deficits. Thus, our results confirm the importance of alteration in cholinergic nuclei in PD and suggest the specificity of the NBM changes in olfactory dysfunction at this early disease stage before the occurrence of cognitive deficits.

#### Contribution of the noradrenergic pathway to olfactory dysfunction in patients with RBD

Neuromelanin signal changes in the LC correlated with UPSIT scores suggesting that this projection was also involved in olfactory deficit in PD. However, reduced noradrenergic receptors-enriched functional connectivity of the NBM was observed with posterior temporo-parieto-occipital regions in patients with RBD only, but not in PD patients without RBD whether anosmic or normosmic. In the single-hit hypothesis of PD^49^, patients with RBD are associated with a vagal origin of the synucleopathy and a “Body-first” progression. Postmortem analysis of patients in early stages of α-synuclein pathology showed that patients with Lewy-bodies constrained in the vagal nerve and locus coeruleus had no pathology in the olfactory bulbs^50^. Olfactory dysfunction in patients with RBD thus seems to arise from a different alteration that may be linked to the LC. The LC is the main source of noradrenergic input to the olfactory bulbs^51,52^.

#### Lack of evidence of dopaminergic contribution to olfactory dysfunction in PD

No correlation was found between UPSIT and putaminal DaT-SBR nor with SN volume or NM contrast, which were associated with motor symptoms as already shown^33^. The literature data is discordant regarding the role of the dopaminergic system in olfactory disorders of the disease, with some studies reporting a correlation between striatal dopaminergic function^16,18^ and UPSIT but others only borderline correlation^15^ or no correlation^17^. Overall, results suggest that cholinergic denervation of the cortex is more robustly associated with hyposmia than nigrostriatal dopaminergic denervation in PD as suggested previously^15^.

## Conclusion

Multiparametric analysis of PD and prodromal patients allowed to identify alterations correlating with RBD and anosmia. Specifically, while dopaminergic dysfunction was common to all PD groups, it didn’t correlate with UPSIT. On the other hand, the grey matter volume in the NBM showed significant correlation with UPSIT in all anosmic patients. Furthermore, the NM-CNR in the LC was also associated with olfactory dysfunction, but only in patients with RBD. These results highlight the role of the degeneration of cholinergic and noradrenergic systems in the olfactory dysfunction in PD and suggest a different mechanism underlying olfactory dysfunction in patients with RBD.

## Supporting information

STROBE Checklist

## Acknowledgements

The authors would like to thank all participants involved in the study.

## Author roles

1) Research project: A. Conception, B. Organization, C. Execution; 2) Statistical Analysis: A. Design, B. Execution, C. Review and Critique; 3) Manuscript: A. Writing of the first draft, B. Review and Critique

J.B.P.: 1A,1B,1C, 2A, 2B, 3A

E.K., A.F., S.O., R.V., S.R., F.X.L., E.M., R.G.: 1C, 2A, 2B

I.A., J.C.C., M.V.: 1B, 2C, 3B

N.P., C.G., S.L.: 1A, 1B, 2A, 2C, 3B

## Bibliography

1. Postuma RB, Berg D, Stern M, et al. MDS clinical diagnostic criteria for Parkinson’s disease. Mov Disord Off J Mov Disord Soc. 2015;30(12):1591–1601. doi:10.1002/mds.26424

2. Postuma RB, Iranzo A, Hu M, et al. Risk and predictors of dementia and parkinsonism in idiopathic REM sleep behaviour disorder: a multicentre study. Brain J Neurol. 2019;142(3):744–759. doi:10.1093/brain/awz030

3. Doty RL. Olfaction in Parkinson’s disease and related disorders. Neurobiol Dis. 2012;46(3):527–552. doi:10.1016/j.nbd.2011.10.026

4. Ubeda-Bañon I, Saiz-Sanchez D, de la Rosa-Prieto C, Martinez-Marcos A. α-Synuclein in the olfactory system in Parkinson’s disease: role of neural connections on spreading pathology. Brain Struct Funct. 2014;219(5):1513–1526. doi:10.1007/s00429-013-0651-2

5. Wang J, You H, Liu JF, Ni DF, Zhang ZX, Guan J. Association of olfactory bulb volume and olfactory sulcus depth with olfactory function in patients with Parkinson disease. AJNR Am J Neuroradiol. 2011;32(4):677–681. doi:10.3174/ajnr.A2350

6. Chen S, Tan H yu, Wu Zhua, et al. Imaging of olfactory bulb and gray matter volumes in brain areas associated with olfactory function in patients with Parkinson’s disease and multiple system atrophy. Eur J Radiol. 2014;83(3):564–570. doi:10.1016/j.ejrad.2013.11.024

7. Sengoku R, Matsushima S, Bono K, et al. Olfactory function combined with morphology distinguishes Parkinson’s disease. Parkinsonism Relat Disord. 2015;21(7):771–777. doi:10.1016/j.parkreldis.2015.05.001

8. Tremblay C, Iravani B, Aubry Lafontaine É, et al. Parkinson’s Disease Affects Functional Connectivity within the Olfactory-Trigeminal Network. J Park Dis. 2020;10(4):1587–1600. doi:10.3233/JPD-202062

9. Dutta D, Karthik K, Holla VV, et al. Olfactory Bulb Volume, Olfactory Sulcus Depth in Parkinson’s Disease, Atypical Parkinsonism. Mov Disord Clin Pract. 2023;10(5):794–801. doi:10.1002/mdc3.13733

10. Wattendorf E, Welge-Lüssen A, Fiedler K, et al. Olfactory impairment predicts brain atrophy in Parkinson’s disease. J Neurosci Off J Soc Neurosci. 2009;29(49):15410–15413. doi:10.1523/JNEUROSCI.1909-09.2009

11. Lee EY, Eslinger PJ, Du G, Kong L, Lewis MM, Huang X. Olfactory-related cortical atrophy is associated with olfactory dysfunction in Parkinson’s disease. Mov Disord Off J Mov Disord Soc. 2014;29(9):1205–1208. doi:10.1002/mds.25829

12. Campabadal A, Uribe C, Segura B, et al. Brain correlates of progressive olfactory loss in Parkinson’s disease. Parkinsonism Relat Disord. 2017;41:44–50. doi:10.1016/j.parkreldis.2017.05.005

13. Barrett MJ, Murphy JM, Zhang J, et al. Olfaction, cholinergic basal forebrain degeneration, and cognition in early Parkinson disease. Parkinsonism Relat Disord. 2021;90:27–32. doi:10.1016/j.parkreldis.2021.07.024

14. Tian Q, An Y, Kitner-Triolo MH, et al. Associations of Olfaction With Longitudinal Trajectories of Brain Volumes and Neuropsychological Function in Older Adults. Neurology. 2023;100(9):e964–e974. doi:10.1212/WNL.0000000000201646

15. Bohnen NI, Müller MLTM, Kotagal V, et al. Olfactory dysfunction, central cholinergic integrity and cognitive impairment in Parkinson’s disease. Brain J Neurol. 2010;133(Pt 6):1747–1754. doi:10.1093/brain/awq079

16. Löhle M, Wolz M, Beuthien-Baumann B, et al. Olfactory dysfunction correlates with putaminal dopamine turnover in early de novo Parkinson’s disease. J Neural Transm Vienna Austria 1996. 2020;127(1):9–16. doi:10.1007/s00702-019-02122-9

17. Goldstein DS, Sewell L, Holmes C. Association of anosmia with autonomic failure in Parkinson disease. Neurology. 2010;74(3):245–251. doi:10.1212/WNL.0b013e3181ca014c

18. Oh YS, Kim JS, Hwang EJ, Lyoo CH. Striatal dopamine uptake and olfactory dysfunction in patients with early Parkinson’s disease. Parkinsonism Relat Disord. 2018;56:47–51. doi:10.1016/j.parkreldis.2018.06.022

19. Meles SK, Vadasz D, Renken RJ, et al. FDG PET, dopamine transporter SPECT, and olfaction: Combining biomarkers in REM sleep behavior disorder. Mov Disord Off J Mov Disord Soc. 2017;32(10):1482–1486. doi:10.1002/mds.27094

20. Chen S, Wang SH, Bai YY, Zhang JW, Zhang HJ. Comparative Study on Topological Properties of the Whole-Brain Functional Connectome in Idiopathic Rapid Eye Movement Sleep Behavior Disorder and Parkinson’s Disease Without RBD. Front Aging Neurosci. 2022;14:820479. doi:10.3389/fnagi.2022.820479

21. Braak H, Tredici KD, Rüb U, de Vos RAI, Jansen Steur ENH, Braak E. Staging of brain pathology related to sporadic Parkinson’s disease. Neurobiol Aging. 2003;24(2):197–211. doi:10.1016/S0197-4580(02)00065-9

22. Luk KC, Kehm V, Carroll J, et al. Pathological α-synuclein transmission initiates Parkinson-like neurodegeneration in nontransgenic mice. Science. 2012;338(6109):949–953. doi:10.1126/science.1227157

23. Horsager J, Andersen KB, Knudsen K, et al. Brain-first versus body-first Parkinson’s disease: a multimodal imaging case-control study. Brain J Neurol. 2020;143(10):3077–3088. doi:10.1093/brain/awaa238

24. Pyatigorskaya N, Yahia-Cherif L, Valabregue R, et al. Parkinson Disease Propagation Using MRI Biomarkers and Partial Least Squares Path Modeling. Neurology. 2021;96(3):e460–e471. doi:10.1212/WNL.0000000000011155

25. Knudsen K, Fedorova TD, Hansen AK, et al. In-vivo staging of pathology in REM sleep behaviour disorder: a multimodality imaging case-control study. Lancet Neurol. 2018;17(7):618–628. doi:10.1016/S1474-4422(18)30162-5

26. Ito E, Inoue Y. [The International Classification of Sleep Disorders, third edition. American Academy of Sleep Medicine. Includes bibliographies and index]. Nihon Rinsho Jpn J Clin Med. 2015;73(6):916–923.

27. Goetz CG, Tilley BC, Shaftman SR, et al. Movement Disorder Society-sponsored revision of the Unified Parkinson’s Disease Rating Scale (MDS-UPDRS): Scale presentation and clinimetric testing results. Mov Disord. 2008;23(15):2129–2170. doi:10.1002/mds.22340

28. Mattis S. Dementia Rating Scale: DRS!ll: Professional Manual. PAR; 1988.

29. Doty RL. Olfactory dysfunction in neurodegenerative diseases: is there a common pathological substrate? Lancet Neurol. 2017;16(6):478–488. doi:10.1016/S1474-4422(17)30123-0

30. Mazziotta JC, Toga AW, Evans A, Fox P, Lancaster J. A Probabilistic Atlas of the Human Brain: Theory and Rationale for Its Development: The International Consortium for Brain Mapping (ICBM). NeuroImage. 1995;2(2, Part A):89–101. doi:10.1006/nimg.1995.1012

31. Mazziotta J, Toga A, Evans A, et al. A Four-Dimensional Probabilistic Atlas of the Human Brain. J Am Med Inform Assoc. 2001;8(5):401–430. doi:10.1136/jamia.2001.0080401

32. Kötter R, Mazziotta J, Toga A, et al. A probabilistic atlas and reference system for the human brain: International Consortium for Brain Mapping (ICBM). Philos Trans R Soc Lond B Biol Sci. 2001;356(1412):1293–1322. doi:10.1098/rstb.2001.0915

33. Biondetti E, Santin MD, Valabrègue R, et al. The spatiotemporal changes in dopamine, neuromelanin and iron characterizing Parkinson’s disease. Brain J Neurol. 2021;144(10):3114–3125. doi:10.1093/brain/awab191

34. Tzourio-Mazoyer N, Landeau B, Papathanassiou D, et al. Automated Anatomical Labeling of Activations in SPM Using a Macroscopic Anatomical Parcellation of the MNI MRI Single-Subject Brain. NeuroImage. 2002;15(1):273–289. doi:10.1006/nimg.2001.0978

35. Yelnik J, Bardinet E, Dormont D, et al. A three-dimensional, histological and deformable atlas of the human basal ganglia. I. Atlas construction based on immunohistochemical and MRI data. NeuroImage. 2007;34(2):618–638. doi:10.1016/j.neuroimage.2006.09.026

36. Bardinet E, Bhattacharjee M, Dormont D, et al. A three-dimensional histological atlas of the human basal ganglia. II. Atlas deformation strategy and evaluation in deep brain stimulation for Parkinson disease: Clinical article. J Neurosurg. 2009;110(2):208–219. doi:10.3171/2008.3.17469

37. Dipasquale O, Selvaggi P, Veronese M, Gabay AS, Turkheimer F, Mehta MA. Receptor-Enriched Analysis of functional connectivity by targets (REACT): A novel, multimodal analytical approach informed by PET to study the pharmacodynamic response of the brain under MDMA. NeuroImage. 2019;195:252–260. doi:10.1016/j.neuroimage.2019.04.007

38. Wu X, Yu C, Fan F, et al. Correlation between progressive changes in piriform cortex and olfactory performance in early Parkinson’s disease. Eur Neurol. 2011;66(2):98–105. doi:10.1159/000329371

39. Stewart SA, Pimer L, Fisk JD, et al. Olfactory Function and Diffusion Tensor Imaging as Markers of Mild Cognitive Impairment in Early Stages of Parkinson’s Disease. Clin EEG Neurosci. 2023;54(1):91–97. doi:10.1177/15500594211058263

40. Nigro P, Chiappiniello A, Simoni S, et al. Changes of olfactory tract in Parkinson’s disease: a DTI tractography study. Neuroradiology. 2021;63(2):235–242. doi:10.1007/s00234-020-02551-4

41. Ibarretxe-Bilbao N, Junque C, Marti MJ, et al. Olfactory impairment in Parkinson’s disease and white matter abnormalities in central olfactory areas: A voxel-based diffusion tensor imaging study. Mov Disord Off J Mov Disord Soc. 2010;25(12):1888–1894. doi:10.1002/mds.23208

42. Ruberg M, Rieger F, Villageois A, Bonnet AM, Agid Y. Acetylcholinesterase and butyrylcholinesterase in frontal cortex and cerebrospinal fluid of demented and non-demented patients with Parkinson’s disease. Brain Res. 1986;362(1):83–91. doi:10.1016/0006-8993(86)91401-0

43. Shimada H, Hirano S, Shinotoh H, et al. Mapping of brain acetylcholinesterase alterations in Lewy body disease by PET. Neurology. 2009;73(4):273–278. doi:10.1212/WNL.0b013e3181ab2b58

44. Liu AKL, Chang RCC, Pearce RKB, Gentleman SM. Nucleus basalis of Meynert revisited: anatomy, history and differential involvement in Alzheimer’s and Parkinson’s disease. Acta Neuropathol (Berl). 2015;129(4):527–540. doi:10.1007/s00401-015-1392-5

45. Mesulam MM. Cholinergic circuitry of the human nucleus basalis and its fate in Alzheimer’s disease. J Comp Neurol. 2013;521(18):4124–4144. doi:10.1002/cne.23415

46. Ray NJ, Bradburn S, Murgatroyd C, et al. In vivo cholinergic basal forebrain atrophy predicts cognitive decline in de novo Parkinson’s disease. Brain J Neurol. 2018;141(1):165–176. doi:10.1093/brain/awx310

47. Schulz J, Pagano G, Fernández Bonfante JA, Wilson H, Politis M. Nucleus basalis of Meynert degeneration precedes and predicts cognitive impairment in Parkinson’s disease. Brain. 2018;141(5):1501–1516. doi:10.1093/brain/awy072

48. Gargouri F, Gallea C, Mongin M, et al. Multimodal magnetic resonance imaging investigation of basal forebrain damage and cognitive deficits in Parkinson’s disease. Mov Disord Off J Mov Disord Soc. 2019;34(4):516–525. doi:10.1002/mds.27561

49. Borghammer P. The α-Synuclein Origin and Connectome Model (SOC Model) of Parkinson’s Disease: Explaining Motor Asymmetry, Non-Motor Phenotypes, and Cognitive Decline. J Park Dis. 11(2):455–474. doi:10.3233/JPD-202481

50. Borghammer P, Just MK, Horsager J, et al. A postmortem study suggests a revision of the dual-hit hypothesis of Parkinson’s disease. Npj Park Dis. 2022;8(1):1–11. doi:10.1038/s41531-022-00436-2

51. Shipley MT, Halloran FJ, de la Torre J. Surprisingly rich projection from locus coeruleus to the olfactory bulb in the rat. Brain Res. 1985;329(1-2):294–299. doi:10.1016/0006-8993(85)90537-2

52. McLean JH, Shipley MT, Nickell WT, Aston-Jones G, Reyher CK. Chemoanatomical organization of the noradrenergic input from locus coeruleus to the olfactory bulb of the adult rat. J Comp Neurol. 1989;285(3):339–349. doi:10.1002/cne.902850305

