## Supplementary Information for "The role of dopaminergic, cholinergic and noradrenergic networks in hyposmia in Parkinson’s disease"

**MRI acquisition**

Whole brain anatomic images were acquired with a T_1_-weighted (T1w) three-dimensional (3D) MP2RAGE sequence (TR/TE=5000/2.98 ms, Resolution = 1x1x1 mm^3^, acquisition time = 08:12 min). NM-sensitive images were acquired using a T_1_w two-dimensional turbo spin echo sequence (TR/TE=890/13 ms, in-plane resolution = 0.3x0.3 mm^2^, 48 slices, slice thickness = 3 mm, acquisition time=06:55) as previously described. Resting state fMRI images were acquired using a single shot gradient echo sequence (TR/TE=1405/30 ms, flip angle 70°, 2.5-mm isovoxel size, 60 interleaved slices, multiband acceleration factor=3, 364 measures, acquisition time=08:41). A blipped acquisition with 6 measures was also acquired for correction of geometric distortions.

**DaTScan acquisition**

Acquisition was performed between 3 and 4h after injection of Lugol regimen for thyroid gland blockage and 185–200 MBq of tracer. Images were acquired with the following parameters: circular orbit, clockwise rotation, zoom factor = 1.5\1.5, matrix size = 128 × 128 voxels, number of slices = 128, voxel size = 2.9×2.9 mm², slice thickness = 2.9 mm, rotations = 1, number of frames in rotation = 120, actual frame duration = 30 s, energy window = 143.1–174.9 keV. Reconstruction was performed on GE Healthcare Xeleris Workstation with iterative algorithm, spatial filtering (fourth-order low-pass filter with a cut-off frequency equal to 3.5 mm^−1^) and Chang attenuation correction (µ = 0.12 cm^−1^).

All the recorded images were visually inspected, and those with severe motion, arterial flow artifacts affecting SN visibility or incomplete SN coverage were excluded.

**Image analysis**

Brain extraction and segmentation into grey matter, white matter and CSF was performed on the whole-brain anatomical MP2RAGE images using Statistical Parametric Mapping software for MATLAB (SPM12). Images were then coregistered to the Montreal Neurological Institute template using nonlinear transformations. These transformations were performed with the Computational Anatomy Toolbox (CAT12) implemented in SPM12 and were used to calculate the deformation needed for each voxel to be registered to the template.

The SN was manually segmented on the NM-sensitive image by two trained raters blind to the groups, as previously described. SN volume was extracted and corrected by total intracranial volume (TIV). Contrast-to-Noise Ratio (NM-CNR) of the SN was computed using a background region including the tegmentum and superior cerebellar peduncle as previously described^37^.

NM-sensitive images of the LC were processed as already described. Briefly, the mean signal of the 10 connected most intense voxels in the LC was automatically computed within large 3D bounding boxes including the LC area. NM-CNR of the LC was defined as the ratio between the average intensity of these 10 brightest connected LC voxels and the mean intensity of a reference region in the rostral pontomesencephalic area averaged over slices to normalize slice intensity.

Cortical areas included the frontal, orbitofrontal, anterior cingulate, temporal, and parahippocampal cortices, as well as the amygdala, insula, and gyrus rectus. Subcortical areas included the caudate nucleus, putamen, thalamus, hippocampus, cerebellum, LC and nucleus basalis of Meynert (NBM). The LC mask and the NBM mask came from previous studies.

**Supplementary Figures**

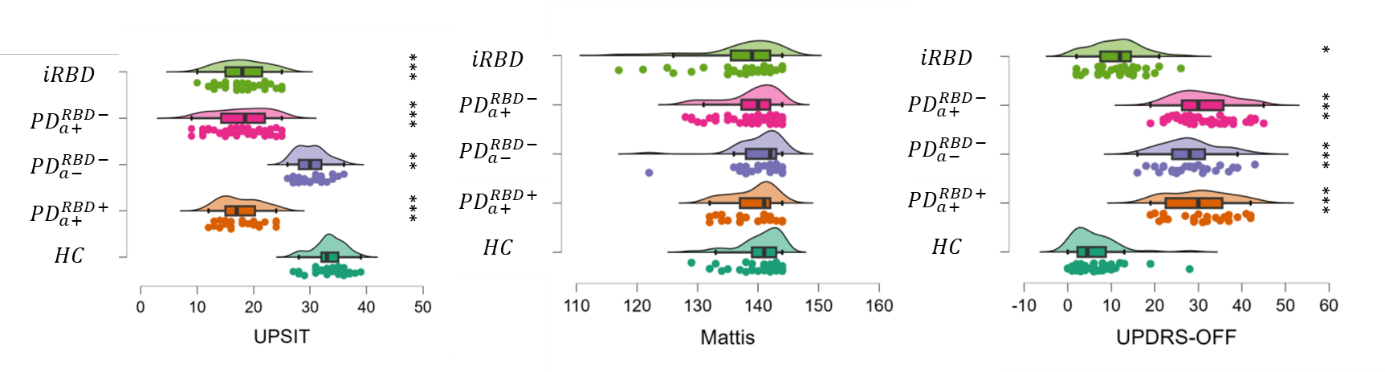

Supplementary Figure 1: UPSIT, Mattis and UPDRS scores of patients. *p<0.05, **p<0.01, ***p<0.001. ANCOVA with age as a covariable followed by Tukey’s post-hoc analysis.

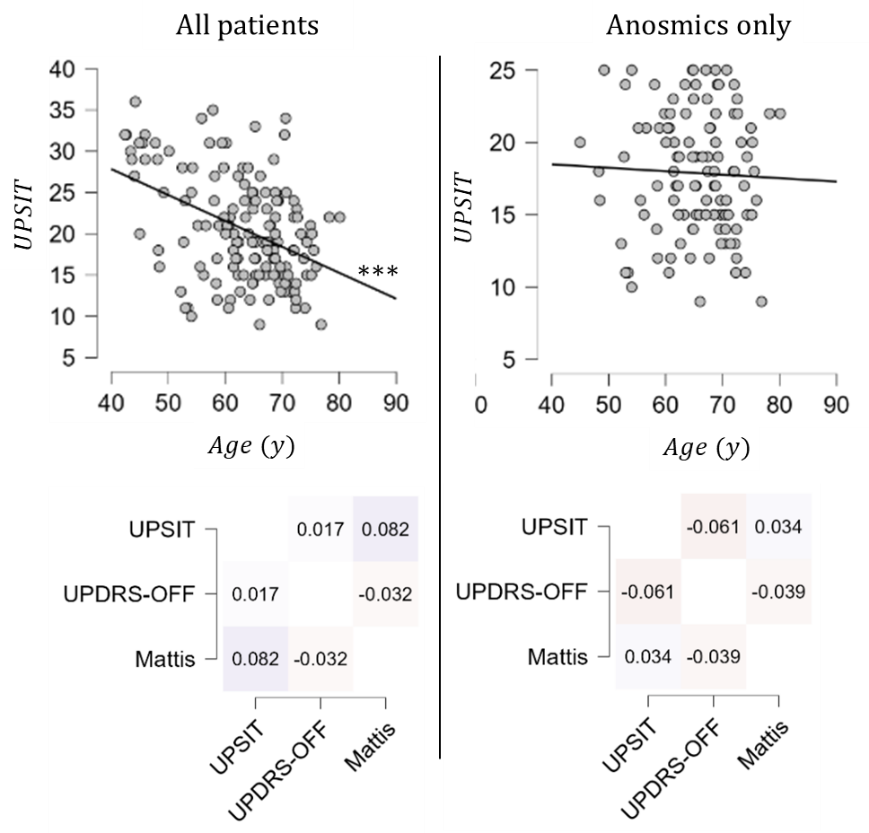

Supplementary Figure 2: Correlations between UPSIT and clinical variables. Left panel: Correlation of UPSIT with age (top line) or Mattis and UPDRS (bottom line) when taking all patients into account. Right panel: Correlation of UPSIT with age (top line) or Mattis and UPDRS (bottom line) when taking only anosmic patients into account. ***p<0.001, Pearson’s Correlation.

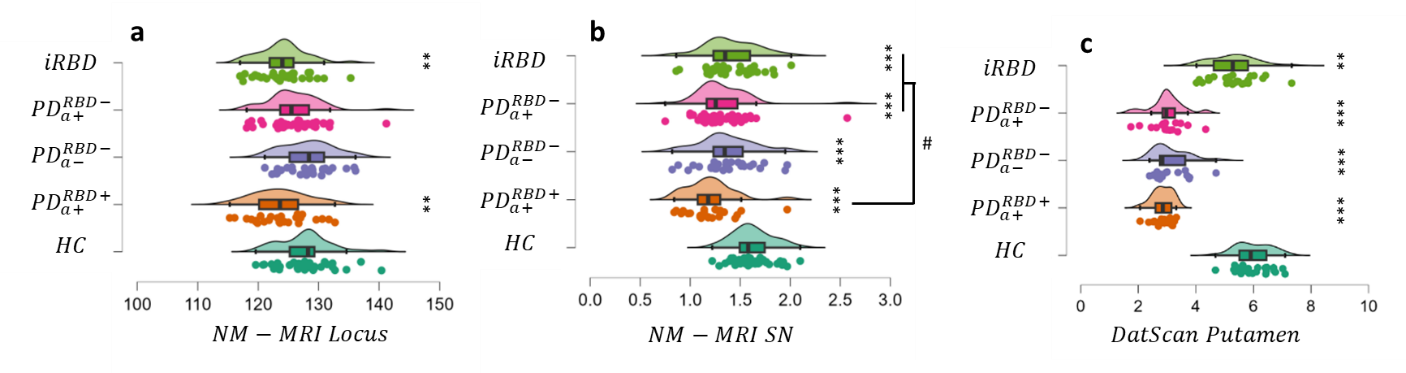

Supplementary figure 3. a) NM-MRI CNR in the locus coeruleus. b) NM-MRI CNR in the substantia nigra. c) DaT striatal binding ratio in the putamen. Significant differences with HC are highlighted *p<0.05, **p<0.01, ***p<0.001. ^#^p<0.05 compared with ${PD}_{a+}^{RBD+}$. ANCOVA with age as a nuisance covariable followed by Tukey’s post hoc.analysis

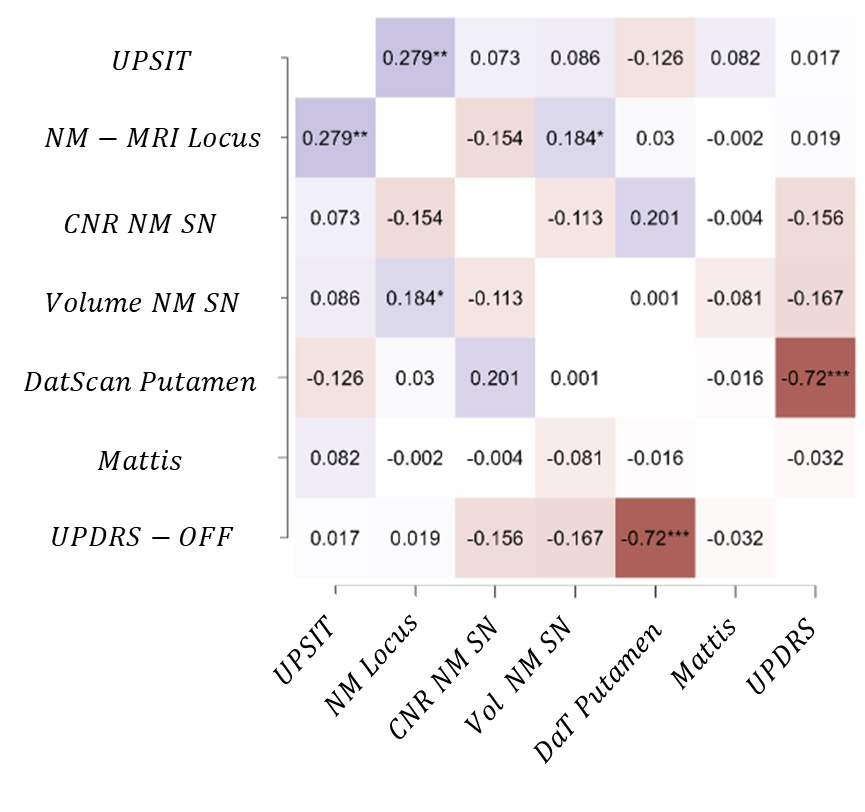

Supplementary Figure 4: Correlation heatmap between all clinical variables, taking all patients into account. *Colors range from dark blue for strong negative correlation to dark red for strong positive correlation* *p<0.05,**p<0.01, ***p<0.001, Pearson’s R correlation coefficient.
